# The Effects of Evolution and Spatial Structure on Diversity in Biological Reserves

**DOI:** 10.1101/043083

**Authors:** Emily L. Dolson, Michael J. Wiser, Charles Ofria

## Abstract

Conservation ecologists have long argued over the best way of placing reserves across an environment to maximize population diversity. Many have studied the effect of protecting many small regions of an ecosystem vs. a single large region, with varied results. However, this research tends to ignore evolutionary dynamics under the rationale that the spatiotemporal scale required is prohibitive. We used the Avida digital evolution research platform to overcome this barrier and study the response of phenotypic diversity to eight different reserve placement configurations. The capacity for mutation,and therefore evolution, substantially altered the dynamics of diversity in the population. When mutations were allowed, reserve configurations involving a greater number of consequently smaller reserves were substantially more effective at maintaining existing diversity and generating new diversity. However, when mutations were disallowed, reserve configuration had little effect on diversity generation and maintenance. While further research is necessary before translating these results into policy decisions, this study demonstrates the importance of considering evolution when making such decisions and suggests that a larger number of smaller reserves may have evolutionary benefits.

## Introduction

Protecting biodiversity is generally acknowledged to be an important conservation goal for a number of reasons, including biodiversity’s role in maintaining various ecosystem services (e.g. carbon sequestration), and its potential as a reservoir of useful and undiscovered genetic innovations (Gaston and Spicer, 2004; Hassan et al., 2005; Loreau et al., 2001; Montoya et al., 2012). However, there is another reason that biodiversity is critically important, which is often overlooked – continued evolution requires diversity. Since adaptation to new environments will be a critical component of the long-term survival of many lineages in the face of climate change, it is important to consider conservation of biodiversity in the context of evolution (Stockwell et al., 2003; Mace and Purvis, 2008; Smith et al., 2014).

Most conservation biology research requires a broad spatial scale. Incorporating the long temporal scale required to study evolution makes this already challenging problem intractable in most cases. As a result, most attempts to factor evolution into conservation planning decisions have, out of necessity, been based on general evolutionary principles rather than empirical analysis of their likely outcomes (Cowling and Pressey, 2001; Sgro et al., 2011; Ferrire et al., 2004). In particular, little research on conservation schemes to date has taken evolution into account. Artificial Life techniques such as digital evolution have a lot of potential as an approach to overcoming these obstacles; they allow for the formation of interesting ecologies in a system with a fast enough generation time to do large scale evolution experiments.

Here, we use digital evolution to revisit the single-large vs. several small (SLOSS) debate from a perspective that incorporates evolutionary theory. The SLOSS debates emerged from the theory of island biogeography (MacArthur and Wilson, 1967). The original argument was that, since large islands have more species, larger reserves should be better for conserving biodiversity (Diamond, 1975). However, this effect is counterbalanced by the fact that placing more reserves might result in sampling from multiple different species pools (Simberloff and Abele, 1976). More recent refinements have considered the placement of the reserves relative to each other and interconnectivity between them (Saunders et al., 1991; Tjorve, 2010).

Evolutionary dynamics likely add additional weight to the argument for several small reserves for a number of reasons. First, transient fitness gains can result in a single lineage sweeping a reserve relatively quickly and wiping out standing diversity. Second, separating reserves decreases the colonization rate, giving other lineages time to gain beneficial mutations of their own (Whitley et al., 1998; Tomassini, 2005). Third, spatial isolation can increase the likelihood of speciation.

Many factors interact to bring about the complex spatial eco-evolutionary dynamics that we observe in biological ecosystems. Indeed, the interactions between various factors are a large part of the reason that the relative benefits of different reserve placement strategies have been so hard to untangle. Here, we seek only to lay the groundwork for addressing the impact of evolution on these questions. In order to facilitate this, we will deal with the simplest possible case: a population of sessile, asexual organisms at the same trophic level. Movement, sexual recombination, and predation likely have dramatic impacts on the resulting dynamics. However, in order to understand these effects, we must first understand the behavior of a system without them. Additionally, for the purposes of this paper, we assume an entirely homogeneous environment, eliminating the possibility for complex interactions among habitat heterogeneity, species diversity, and reserve area (Kadmon and Allouche, 2007).

## Methods

### Study System

We conducted our experiments *in silico*, using the Avida Digital Evolution Platform version 2.12.4 (available at https://www.github.com/devosoft/avida) (Ofria and Wilke, 2004). The world of Avida is a two-dimensional grid of cells occupied by digital organisms. These organisms are actually computer programs; their genomes are sequences of simple computer instructions. At the beginning of the experiment, we seed the world with a single ancestor that contains the instructions necessary to copy itself. As organisms copy themselves, they periodically make mistakes, introducing mutations. Some of these mutations will improve the efficiency of self-replication, so the organisms that have them will copy themselves faster than the others. If there is no space available for an organism’s offspring, the offspring will replace an existing organism. As a result, there is selection for organisms that can replicate themselves faster. Because there are mutation, inheritance, and selection, evolution by natural selection occurs.

To allow for the formation of more complex ecologies, we can also choose to reward organisms for performing various computational tasks by allowing them to execute their genomes faster. These tasks can be thought of as pathways for metabolizing various resources, and allow for different organisms to specialize on different survival strategies. To allow for the formation of a stable ecosystem, we can establish negative frequency dependence by linking each task to a limited resource, such that organisms are rewarded for a task in proportion to the amount of the relevant resource that they have access to (Chow et al., 2004).

### Experimental Set-up

For this study, we started by evolving ten populations in the limited resource environment described above (Chow et al., 2004). Each population was started from the same hand-coded self-replicator, but was then allowed to diverge for 100,000 updates, a length of time roughly equivalent to 2000 generations. We then placed these populations in one of eight environments for 100,000 more updates. Each environment had reserves placed across it in a different configuration (see Figure 1). All environments had a total of 900 out of the 3600 grid cells placed in square reserves that tiled evenly across the environment. Reserve configurations varied from having many very small reserves (900 1 by 1 reserves at the extreme) to having a single very large reserve (one 30 by 30 reserve at the extreme). Because world size was held constant, configurations with more reserves necessarily involved them being placed closer together (there was always one reserve worth of space between reserves on all sides). Organisms living in areas outside of the reserves were at risk of being randomly killed each update, according to a specified kill rate (100, 200, 300, 400, or 500 cells were selected per update). Offspring were placed probabilistically near their parents, according to a Poisson distribution, to create a spatial population structure roughly analogous to tree seed dispersal. In order to ascertain what effect allowing populations to evolve was having on our results, we also ran a series of controls in which mutations were disallowed for the second 100,000 updates. We ran 10 replicates per treatment in a fully factorial design across initial population, environment, five kill rates, and mutations being allowed vs. disallowed.

**Figure 1:**
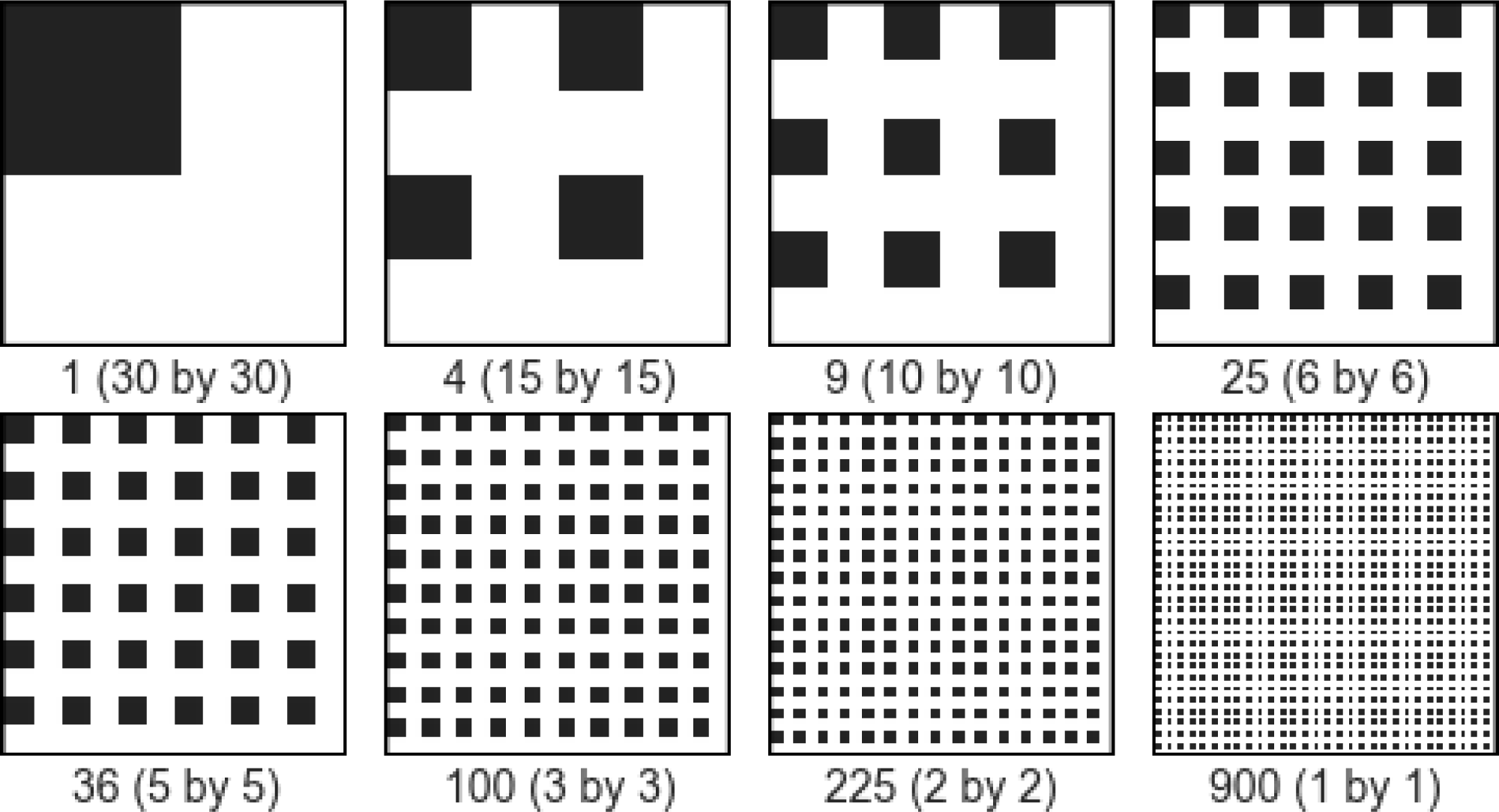
Reserve configurations. Black cells are part of reserves, while white cells are unprotected. Note that the world is toroidal, so spacing between all reserves within a condition is equivalent.

### Analysis

There are three possible mechanisms by which our reserve schemes could drive changes in diversity over evolutionary time - a reserve configuration might: 1) sample from a different range of locations across the environment, 2) promote improved maintenance of existing diversity, and/or 3) promote improved generation of new diversity. To address possible mechanism (1), we measured the number of phenotypes in any reserve at the beginning of the experiment for each condition. To address possible mechanisms (2) and (3), we collected a variety of data on the phenotype-area relationships in our data: phenotype richness within each reserve, total phenotype richness captured across all reserves in a replicate, count of phenotypes lost over the experimental treatment, and count of new phenotypes that evolved over the course of the experimental treatment.

All analyses were conducted using the R Statistical Computing Language, version 3.2.3 (Team, 2013). For statistics in which the unit of replication was a single run of Avida (phenotype loss and generation of novelty), we used a 2-way ANOVA in which initial population, reserve size, and their interaction were treated as random effects. Effect sizes were calculated as partial eta squared (Lakens, 2013). We also calculated some statistics (alpha richness) for which the unit of replication was individual reserves. To account for the non-independence that this set-up introduced, we used linear mixed models with random effects for initial population and replicate, as implemented in the lme4 R package (Bates et al., 2015).

## Results and Discussion

Overall, the replicates in which mutations were allowed during the second 100,000 updates had substantially higher diversity (both richness and Shannon entropy) at the end of the experiment than replicates for which mutations were disallowed (see Figure 3). Runs in which mutations were disallowed had an average of 192.009 +‐ 1.345 fewer phenotypes remaining at the end than runs in which evolution was allowed to continue (Linear mixed model, Chi-squared = 10123, p <.0001). Most ecological models do not include this drop-off (MacArthur and Wilson, 1967; Kadmon and Allouche, 2007; Tjorve, 2010). This discrepancy is partially because ecological models of reserve placement are based on models of island biogeography, and so do not include a phase prior to reserve placement. However, this is a superficial distinction. The more fundamental explanation is likely that most ecological models are built on the assumption of some sort of competitive equivalency between phenotypes, usually based off of the idea that all extant phenotypes are well-optimized to their environment. We make no such assumption. Instead, diversity in these experiments is stabilized through negative frequency dependence (due to limited resources, discussed above) and co-evolutionary arms races, resulting in a dynamic near-equilibrium. While these mechanisms are more realistic, they generally mean that advantages that one lineage has over another are unlikely to persist in the long-term. At any arbitrary point in time, there is probably a lineage with a slight advantage over other lineages. Removing mutations will eliminate the generation of novelty and leave this lineage at an advantage for the rest of the experiment.

Allowing ongoing mutations dramatically increased the extent to which a greater number of smaller reserves promote higher final phenotypic richness than configurations with a smaller number of larger reserves (see Figure 3). For the runs in which mutations were allowed during the second half, increasing the log of the number of reserves by one increased the log of the final number of phenotypes by approximately .0956 +‐ .00259 (Linear mixed model, Chi-squared = 1172.6, p <.0001). This effect was substantially weaker among runs where mutations were disallowed, although still significantly different from 0 (Linear mixed model, Chi-squared = 25.298, p <.0001, slope=0.0084 +‐ 0.0017 log(phenotype count) per additional log(reserve count) units). This appears to be the result of a combination of the three mechanisms described in the Methods section.

### More/smaller reserves capture more phenotypes

Configurations with a greater number of consequently smaller reserves captured a greater number of phenotypes in reserves, likely due to the substantial clumping of closely related organisms (high spatial autocorrelation). In the ten initial populations, the relationship between phenotype richness within a reserve and the size of that reserve followed the pattern of a standard species-area relationship (see Figure 2), with a positive linear relationship between the logarithms of reserve size and reserve richness (Connor and McCoy, 1979). Despite this positive relationship, the total phenotypic richness summed across all reserves within an initial environment was negatively correlated with the area of each of those reserves. This negative relationship strengthened when populations were allowed to evolve for 100,000 updates in the reserve design, while the slope of the positive relationship between within-reserve richness and reserve area decreased slightly (see Figure 3). These effects both weakened dramatically when mutations were disallowed, but remained significantly different from zero. This ability for many small reserves to sample across multiple species pools is precisely the scenario that Simberloff and Abele initially brought up as a counterexample to the argument that a smaller number of larger reserves was always preferable (Simberloff and Abele, 1976). Because organisms disperse locally, similar phenotypes are likely to be clumped together in space, effectively creating multiple species pools.

**Figure 2:**
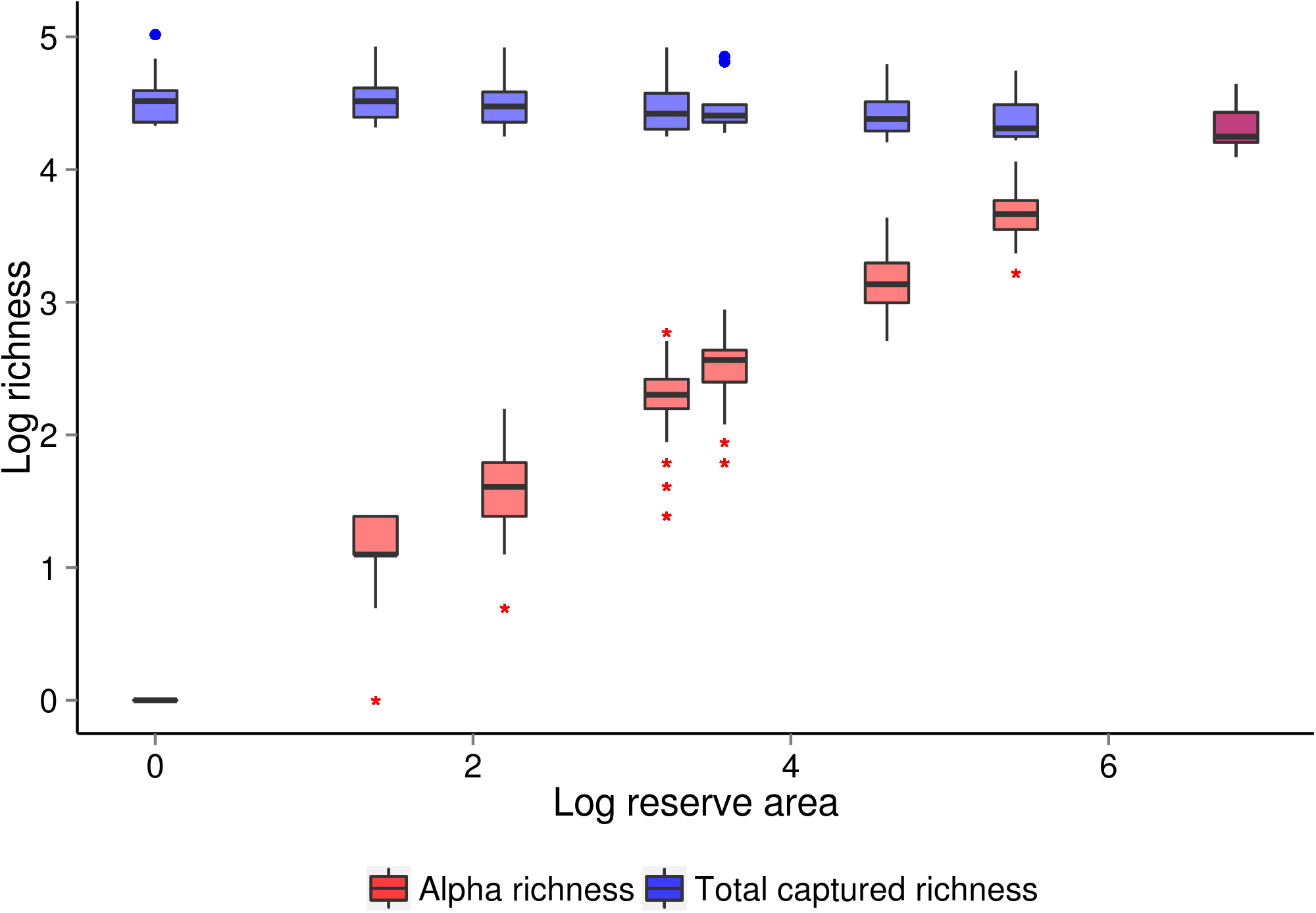
Richness captured within and across reserves at the start of the experiment. Red boxes show the log of the number of phenotypes captured within each reserve across all runs. Blue boxes show the log of the total number of phenotypes captured across all reserves. Note that, despite the clear positive relationship between the size of a reserve and its phenotypic richness, the highest total richness across reserves is achieved in configurations with small reserves.

**Figure 3.**
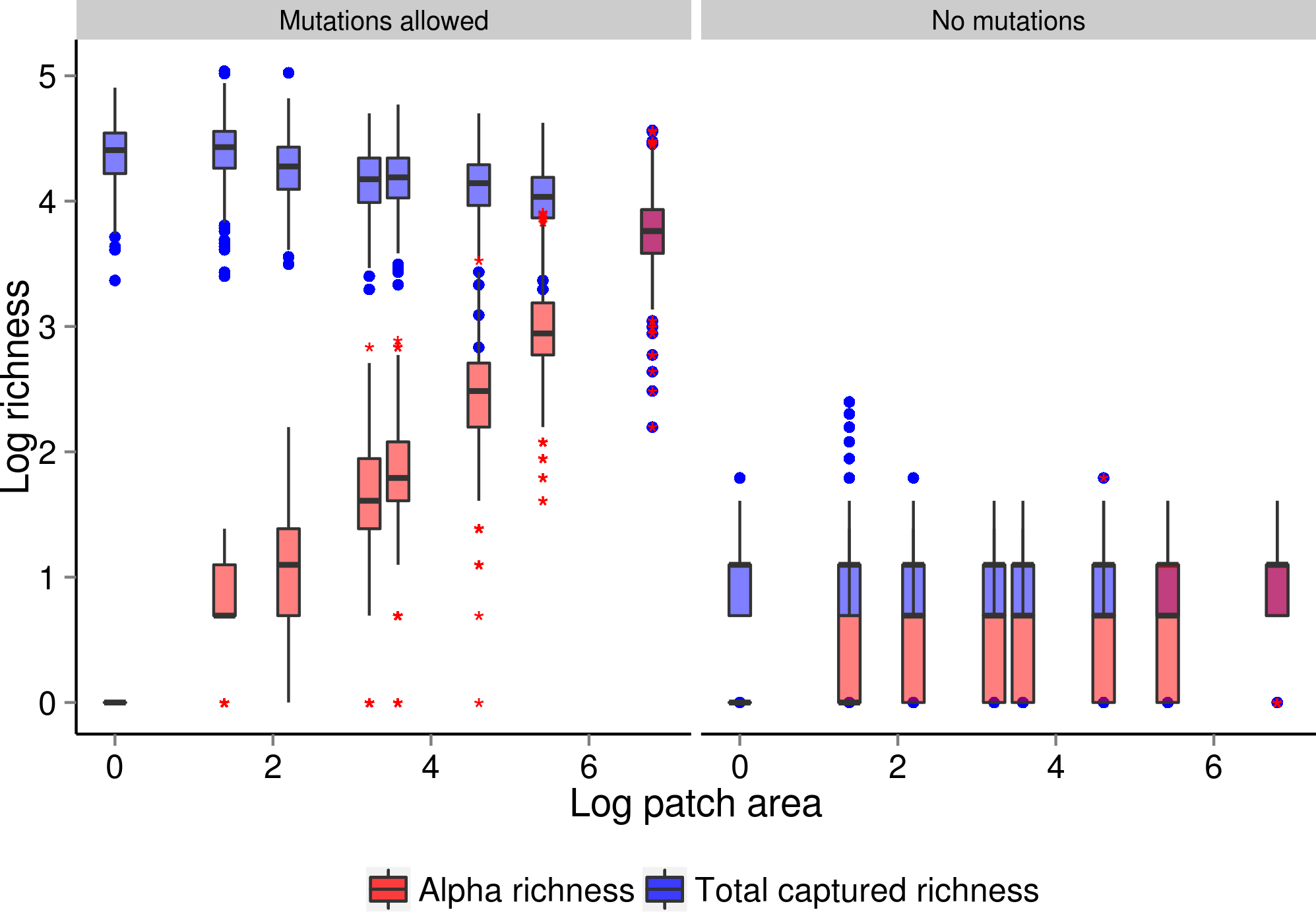
Richness within and across reserves at the end of the experiment for both conditions. Note that the negative relationship between reserve area and total captured richness has intensified since the beginning of the experiment for runs in which mutations are allowed.

### More/smaller reserves promote diversity maintenance

We measured the number of phenotypes that were present in the initial population but no longer present in the final population, i.e. lost phenotypes (see Figure 4). When mutations were allowed, there was a strong positive relationship between phenotype loss and reserve sizes - environments with larger reserves resulted in greater phenotype extinction by the end of the experiment (two-way ANOVA, F(1,3980) = 1680.26, p <.0001, 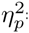=.296). Among the runs where mutations were disallowed, however, there was no significant effect of reserve configuration on phenotype loss (two-way ANOVA, F(1, 3980) = 1.17, p = .41). This discrepancy is an example of the vast impact that allowing for evolutionary dynamics can have.

**Figure 4:**
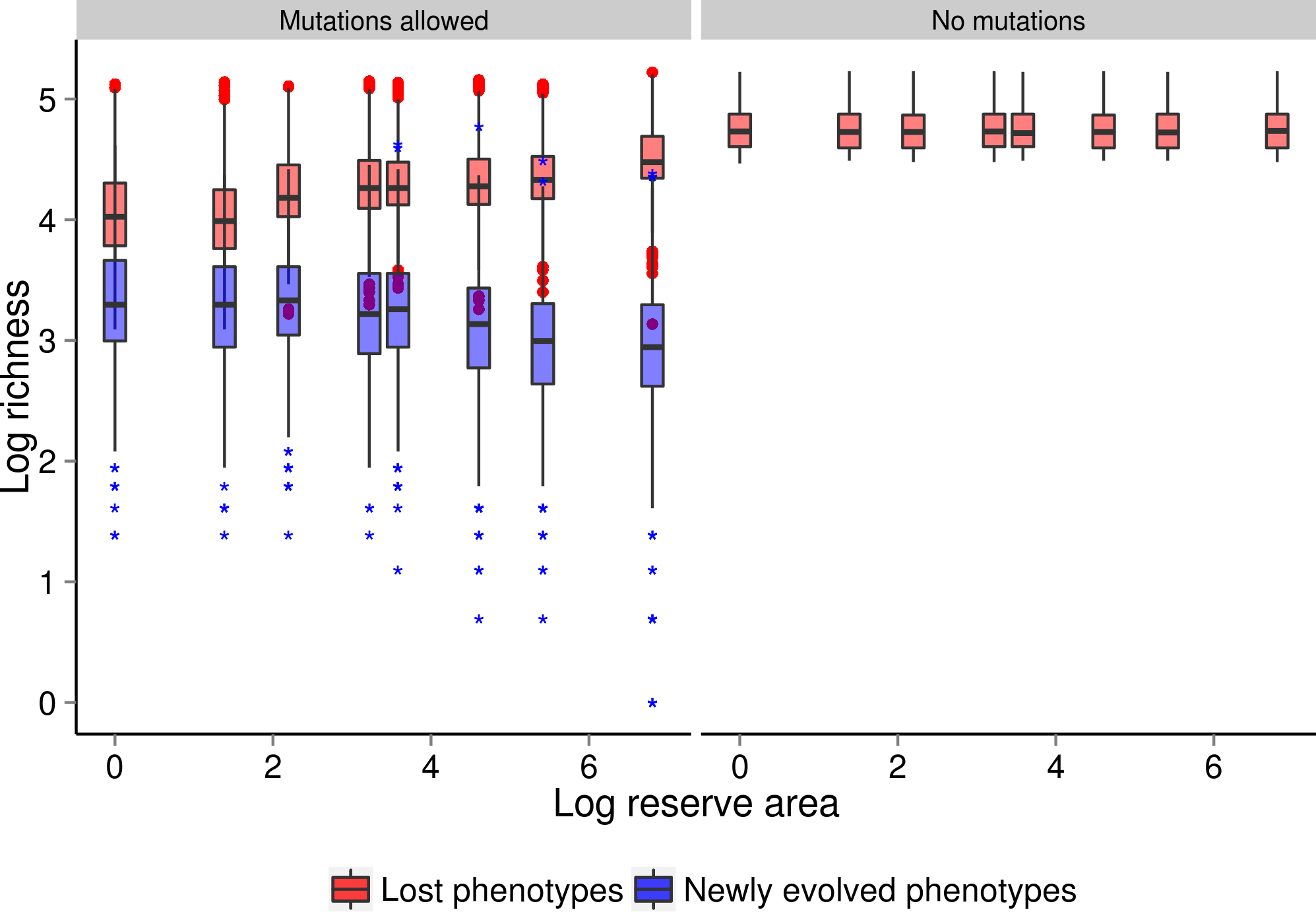
Diversity maintenance and generation across conditions. Red boxes show the log of the number of phenotypes that were initially captured in reserves but were not present at the end of the experiment. Blue boxes show the log of the number of phenotypes that were not initially captured in reserves but were present at the end of the experiment. Note the positive correlation between phenotype loss and reserve size when mutations are allowed.

There are a number of potential evolutionary drivers behind this effect, mostly related to the dynamics of selective sweeps in the population. In reproducing populations, some individuals will have more offspring than others. When one genotype has a substantial selective advantage over its competitors, selection will favor a rapid increase in that genotype’s relative frequency, driving competitors in the population toward extinction. Such a process is referred to a selective sweep, as selection sweeps less-fit variants out of the population McVean (2007). Selective sweeps typically dramatically reduce diversity within the affected population. Not only will the region of the genome under selection go to fixation in the population, but other variant sites on the genetic background where the beneficial trait first arose may also fix. Selective sweeps may be incomplete if, for example, a lineage encounters a competitor with too similar of fitness. An incomplete selective sweep may also occur if a competitor exhibits negative frequency dependence and thus reaches an equilibrium. Even incomplete selective sweeps can substantially reduce diversity, by making a large fraction of the population identical, and driving competing variants extinct (Biswas and Akey, 2006).

Selective sweeps occur faster in populations with higher strength of selection and spatial connectivity (Cantu-Paz, 2001). Both of these factors are impacted by reserve placement; large unprotected regions that fall between reserves decrease connectivity between those reserves (see Figure 5), effectively breaking them into multiple subpopulations and thereby decreasing strength of selection (Gavrilets and Vose, 2005). In this experiment, inter-reserve connectivity and reserve size are inversely correlated because we are holding the size of the entire world constant. As a result, it is hard to determine the impact of each of these variables on the dynamics of selective sweeps in this experiment. Based on the empirical results, however, the effect of reduced subpopulation size would appear to be stronger than the effect of increased connectivity.

The initial population had a significant effect on diversity maintenance, both when mutations were allowed (two-way ANOVA, F(9, 3980) = 1602.45, p <.0001) and disallowed (two-way ANOVA, F(9, 3980) = 204019.41, p <.0001). When mutations were allowed, there was also a significant interaction between reserve configuration and initial population (two-way ANOVA, F(9, 3980) = 7.99, p <.0001). The interaction term is likely significant because some reserve configurations happen to capture more of the initial diversity of any given initial population than others. There may also be an effect of some initial populations having higher starting diversity, stability against selective sweeps, or evolutionary potential than others.

### More/smaller reserves promote diversity generation

We measured the number of phenotypes that were not present in the initial population but were present in the final population, i.e. newly evolved phenotypes (see Figure 4). Among the runs where mutations were allowed, the count of newly evolved phenotypes had a negative relationship with reserve size - environments with larger reserves resulted in fewer newly evolved phenotypes in total (two-way ANOVA, F(1, 3980) = 278.47, p <.0001, 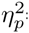=.065). This relation-ship suggests that smaller reserves do a better job of generating new diversity. While the effect is much weaker than the effect of smaller reserves on diversity maintenance, it is still substantial (Lakens, 2013). As with diversity maintenance, it is likely that this effect is driven by dynamics related to selective sweeps. When organisms are competing against fewer other organisms for space, as is the case in a smaller reserve (but see Figure 5), the strength of selection is weakened. This is generally believed to allow time for evolutionary innovation and increased diversification into new niches. A configuration with many small reserves can be thought of as roughly equivalent to an evolutionary algorithm that maintains multiple sub-populations and allows occasional migration between them. Such algorithms are generally quite efficient, perhaps due to their improved ability to maintain and generate diversity (Tomassini, 2005).

**Figure 5.**
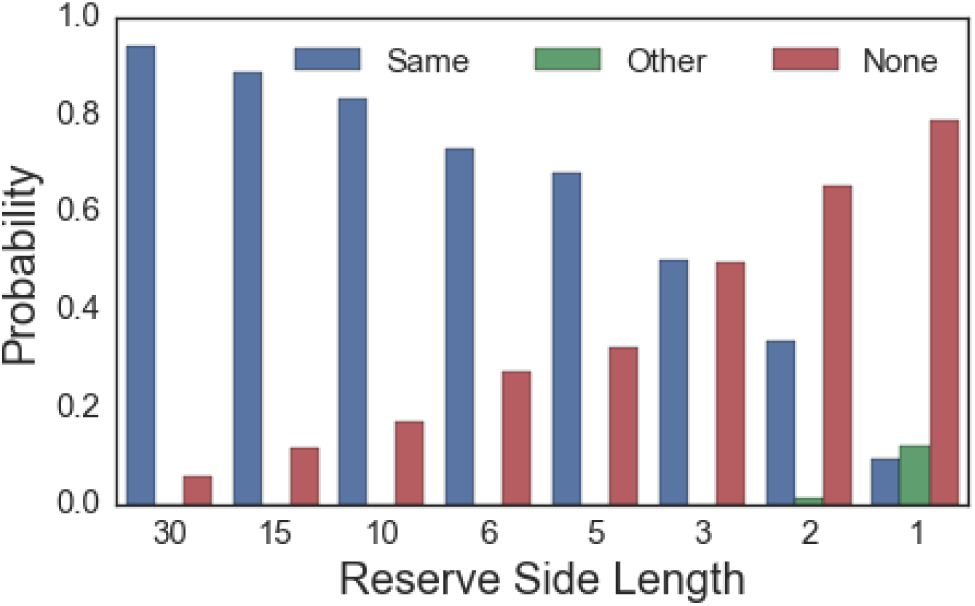
Reserve connectivity increases as patch size decreases. Bars show percentage of offspring from an arbitrary focal reserve that end up in the same reserve (blue), a different reserve (green), and no reserve (red). Note that reproduction events where a parent from one reserve has offspring in another reserve are incredibly rare in all reserve configurations except the two in which reserves are smallest.

## Conclusions

We have laid groundwork for an integration of evolutionary dynamics into reserve design. Our results suggest that evolution fundamentally changes the way that reserve placement affects diversity. In the presence of evolution, configurations with a greater number of consequently smaller reserves substantially improved diversity maintenance. However, in the absence of evolution, there was no effect of reserve configuration on maintenance of the captured initial diversity. Configurations with a greater number of smaller reserves also promoted the evolution of a larger number of novel phenotypes, a dynamic that is only possible when mutations are allowed. These results extend prior research on island model genetic algorithms to address a multi-niche ecosystem, which provides abetter analog to biological ecosystems where conservation applications are relevant.

While our results have potentially important implications for conservation management decisions, it is important to recognize that a number of our simplifying assumptions will bias our results against a smaller number of larger reserves. None of the organisms in our experiment are, for instance, larger or at a higher trophic level than any other organisms. In nature, some organisms require vastly more space than others in order to have access to sufficient energy sources. Additionally, as all of our organisms are sessile, they are not at risk of wandering out of small reserves, as mobile organisms would be. Similarly, as the organisms considered here are asexual, factors such as inbreeding depression and Allee effects are not accounted for. This research represents the first step in an understanding of how evolution interacts with reserve placement. Other topics for future research include the effect of corridors, distance between reserves, placement of reserves in relation to spatial resources, interactions with motile organisms, and the impact of sexual recombination and gene flow on diversity in these systems.

## Acknowledgements

We extend our thanks to Phoebe Zarnetske for her guidance on the ecological theory and spatial analysis in this paper. This research has been supported in part by the National Science Foundation (NSF) BEACON Center under Cooperative Agreement DBI-0939454, by the National Science Foundation Graduate Research Fellowship under Grant No. DGE-1424871, and by Michigan State University through computational resources provided by the Institute for Cyber-Enabled Research. Any opinions, findings, and conclusions or recommendations expressed in this material are those of the author(s) and do not necessarily reflect the views of the NSF.

